# Temporal expression divergence of network modules

**DOI:** 10.1101/167734

**Authors:** Yongjin Park, Tae-Hyuk Kang, Theodore Friedmann, Joel S. Bader

## Abstract

Here we propose new module-based approaches to identify differentially regulated network sub-modules combining temporal trajectories of expression profiles with static network skeletons. Starting from modules identified by network clustering of static networks, our analysis refines pre-defined genesets by partitioning them into smaller homogeneous sets by non-paramettric Bayesian methods. Especially for case-control time series data we developed multi-time point discriminative models and identified each network module as a mixture or admixture of dynamic discriminative functions. Our results shows that our proposed approach outperformed existing geneset enrichment methods in simulation studies. Moreover we applied the methods to neural stem cell differentiation data, and discovered novel modules differentially perturbed in different developmental stages.

## Introduction

### Dynamics of biological networks

Studies on dynamic networks have heavily focused on node-level analysis. Hub nodes of physical interaction and signaling networks were classified to so called “party” and “date” hubs judged by average correlation with neighboring nodes [1]. Despite criticisms [2–4], dynamic property of network and co-expression modules have provided new biological insights into dynamics and systems (e.g., [1, 5–8]). Nevertheless, there still remains subtlety in the analysis. As pointed out [8], node-based statistics and notion of modular structure can be highly data-dependent, but there is no room for heterogeneity in models.

We propose new module-based approaches to dynamic network analysis. However, our notion of module is based on network topology, not expressions. These structural modules can be inferred from static edges. Previously we estimated a trace of dynamic modules while perturbing edges by expressions [9]. Although modules are usually fixed and static, we strive to relax other aspects. We do not attempt to summarize overall node-node relations by a single metric. Network modules are not fixed, but properties of contained nodes change over time. Moreover, we can circumvent long-standing “*n* ≪ *p* problem” by not collecting statistics at node or node-pair level. Instead, we assume genes within well-defined modules are independent identically distributed; then, our *n* is not number of samples but number of genes. We turned the problem to opposite *n ≫ p* regime.

In essence our proposed method resembles geneset analysis methods. We use interaction maps and expression matrix. First we resolve modules from static interactions; then, search for significant dynamic modules by set-based datamining. However, we greatly relaxed assumptions on the sets. Genes are not always identical, i.e., there can be multiple subgroups within genesets. Samples are time-dependent and dynamic. More importantly our goal is to identify significant module at specific timepoints. Modules are not necessarily responsive to environments.

### Time-specific discriminative modules

Regarding gene-wise set-heterogeneity, our solution is quite straightforward. We sought to refine pre-defined genesets by partitioning them into smaller homogeneous sets. We used Bayesian non-parametric method, Dirichlet Process Mixture [10]. However, we need a new paradigm of set-based significance testing to account for sample heterogeneity, especially with time-series samples. Let us put geneset analysis in a classification setting.

Suppose we have a time-series gene expression matrix *X*. Each *x*_*it*_ element measures gene *i*’s mRNA concentration at time *t*. There are *n* genes and *T* time points. Within this dataset, we may assume genes are independent and identically distributed. We also have each gene *i* labeled with *l*_*i*_to indicate that was collected in wild-type (WT) cells if *l*_*i*_ = 1, or abnormal cells if *l_i_* = 0. Then we can fit a logistic regression model with parameter *β_t_* for each time point to classify the gene labels {*l_i_*}, i.e.,

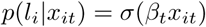

where the sigmoid function 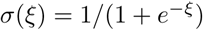. We may estimate *β* by maximizing likelihood function

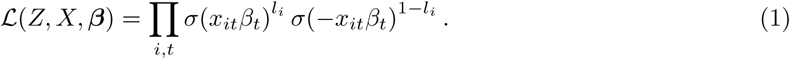

### Temporal Expression Divergence (TED) model

We think of a case-vs-control study in the same classification context. Usually we measure two types of *n × T* gene expression matrices, *X* and *Y* for case and control, or vice versa. Again, we may assume genes are identical and independent, but later we will relax this assumption using predefined genesets. Although the label variable *l*_*i*_not so explicit, we can set all *x*_*it*_ are labeled with *l*_*i*_ = 1 but *y*_*it*_ with *l_i_* = 0. Then, we want to estimate *T* logistic regressors, each parameterized by *β_t_*, maximizing

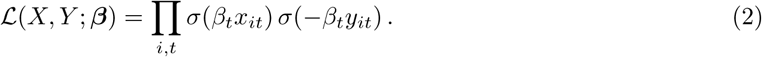

By this model, we can quantify discriminative power of homogeneous *X, Y* time-specifically by magnitude of *β_t_*. Suppose a pair (*x*_*it*_, *y*_*it*_) is discriminative, e.g., *x*_*it*_ ≫ 0 while *y*_*it*_ ≪ 0. Then we maximize the product of two sigmoids with strongly positive *β_t_*. However, if we have both *x*_*it*_ ≫ 0 and *y*_*it*_ ≫ 0, then we want a neutral *β_t_* that is close to 0 to avoid *σ*(−*∞*) = 0.

Fig.1 demonstrates strength of this framework. We generated control datapoints x_*i*_ ~ *N*(x_*i*_|*μ*_case_, *σ*^2^) (colored red) and case y_*i*_ ~ *N*(y_*i*_|*μ*_control_, *σ*^2^) (colored blue). We tested three toy examples generated under different variance parameters *σ* ∈ {0.1,0.5, 0.75} (denoted on the top of subplots). Under small variance *σ* = 0.1, patterns of diverging time points are visually evident. With increasing variance to *σ* = 0.5, datasets became fuzzier and harder to separate out, but the beta parameters, corresponding to decision boundary of time-specific logistic regressors, maintain large values deviating from zero. However, with too large variance *σ* ≥ 0.75, two types of datasets became really indistinguishable, then decision boundary was pulled toward zero. This aspect is also desirable. We want to say “I don’t know” for unclear case-versus-control.

**Figure 1.**
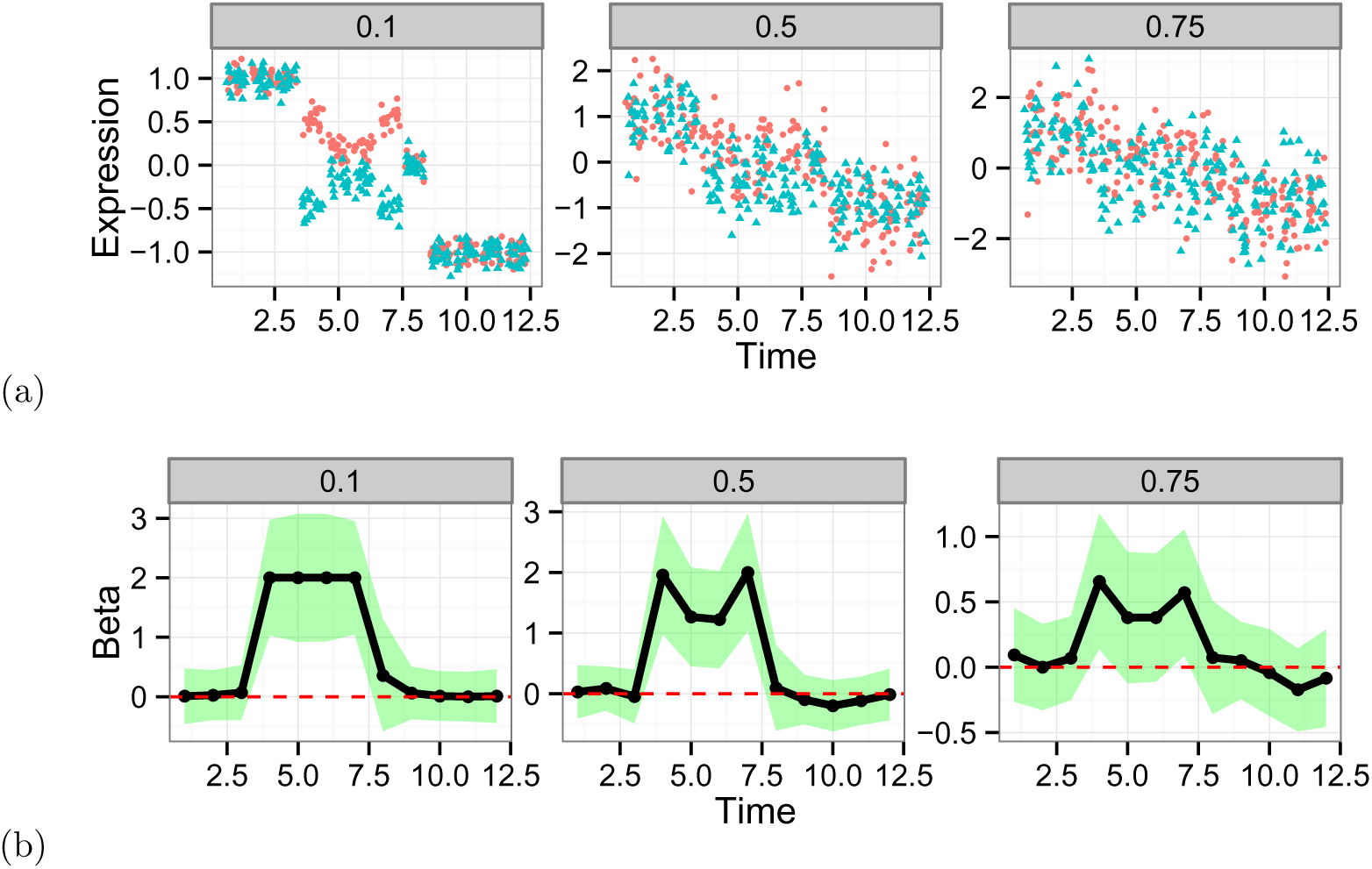
Demonstration of Temporal Expression Divergence on datasets with time-dependent batch effects. (a) We simulated datasets varying standard deviation parameter *σ* = 0.1, 0.5, 0.75 while fixing mean vectors for case (red) and control (blue) samples to *μ*_case_ = [1, 1, 1, .5, .2, .2, .5, 0, −1, −1, −1, −1] and *μ*_control_ = [1, 1, 1, −.5, −.2, −.2, −.5, 0, −1, −1, −1, −1]. For each *σ*, we generated 20 of x_*i*_ ~ *N*(x_*i*_|*μ*_case_, *σ*^2^*I*) and y_*i*_ ~ *N*(y_*i*_|*μ*_control_, *σ*^2^*I*). (b) TED model parameters *β* fitted by our Bayesian inference algorithm (see Methods). Think solid black lines indicate 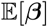 and shaded areas show confidence bands within 2 standard deviation.

### Contributions

Our technical contributions are in two folds: (1) we developed highly sensitive and accurate discriminative model for geneset analysis; (2) we then applied TED models in two well known modeling frameworks, Dirichlet Process Mixture [10] and Latent Dirichlet Allocation [11]. Moreover, from systematic analysis we were able to propose new hypothesis of disease mechanism of neural disorder; and we revealed systematic bias of set- or pathway-based analysis and discussed new approaches.

## Results and Discussion

### Comparison with other geneset analysis methods

We compared performance of TED and TED-dpm with existing methods. Although exhaustive comparison with all existing methods could be beneficial, we chose: generally applicable geneset enrichment with full pairwise sample comparison (GAGE), GAGE with paired sample comparison (GAGE-paired) [13], and geneset analysis (GSA) [14]. We chose GAGE over PAGE [15] because of better performance [13]; GSA over others because of generally good performance in empirical comparison [16]. We did not include the work of Goeman [17] based upon generalized linear models (GLMs) [18] fitting since our TED model is also a special case of generalized linear model with temporal specificity.

Two types of setwise expression matrices of 10 samples were generated; one under the null hypothesis (marked *H*_0_) and alternative (*H*_1_). Under the null hypothesis, both control *x_it_* and case *y_it_* expressions were sampled from the standard Normal distribution, i.e., *x_it_*|*H*_0_ ~ *N*(*x_it_*|0,1) for all *t* ∈ [10]. However, whiting genesets under the alternative hypothesis, 60% of genes were differentially expressed; *x_it_* ~ *N*(*x_it_*|−.5,1) while y_it_ ~ *N*(*y_it_*|-.5,1). All the method performed similarly well on completely differential expression (100%). In addition, we tested conditions where not all time points were differntially expressed under the alternative. We included a certain fraction of non-informative time points, following the null distribution to the differentiating genesets, from 0% to 60%.

Fig.2 summarizes the results. Each column show dispersion of adjusted *p*-values found by different methods under the null (*H*_0_) and alternative (*H*_1_) hypotheses. We expect an oracle method, which gives perfect separation, would generate *p*-values clustered to 1 under the null, but 0 under the alternative. Without noisy samples (the 1st column), all the methods work well as expected. However, as we introduce noisy samples more, we may not pick up significant fraction of true positives using GAGE or GSA. Especially when 60 % noisy samples introduced, GAGE and GSA become significantly underpowered compared to TED. However, TED, not using DPM-based set refinement, might have higher false discovery rate than other methods (outliers of the boxplots); yet, the set refinement method corrected this adequately.

**Figure 2.**
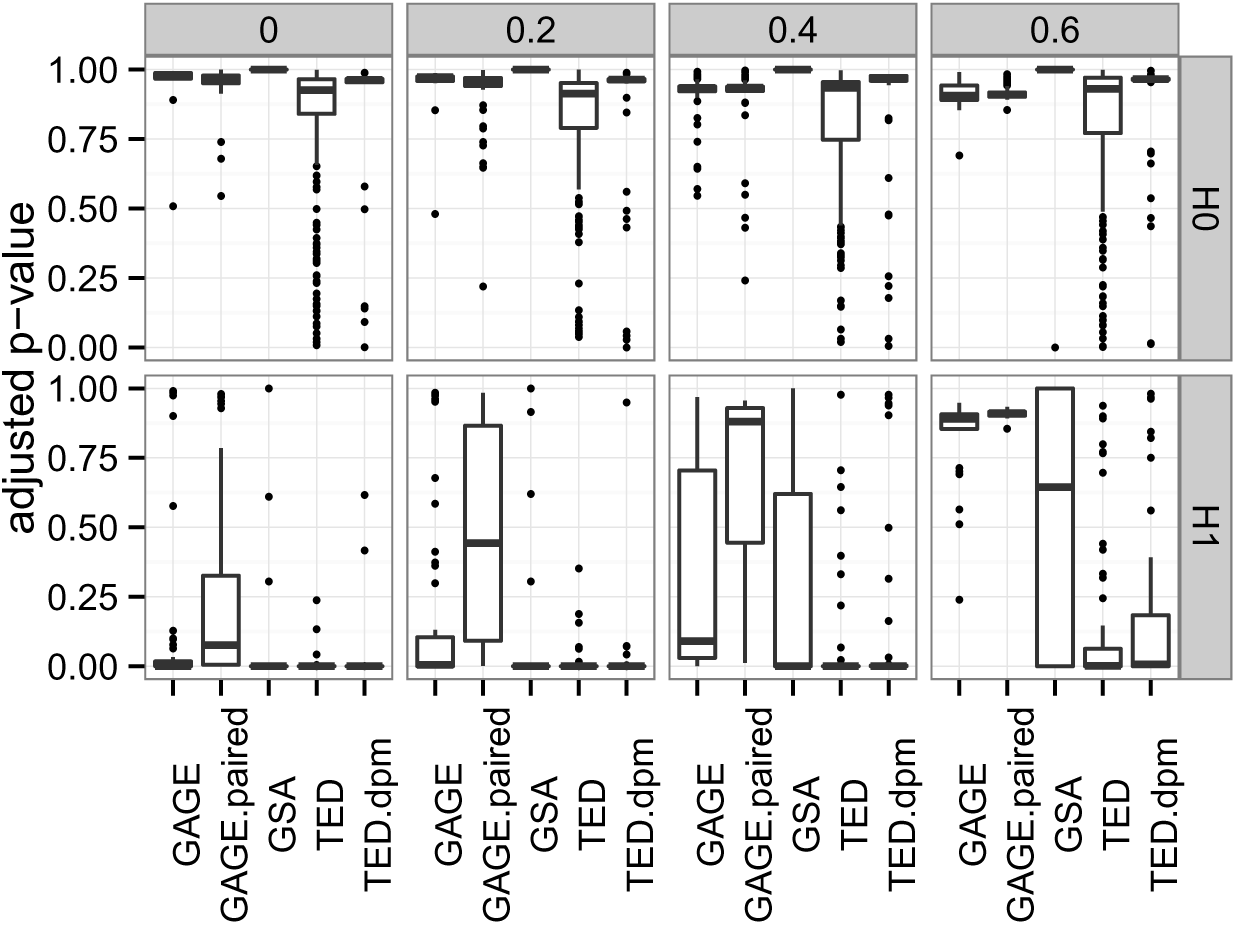
Performance of geneset enrichment analysis. Abbreviation of methods are stated in the text. Each method reported *p*-values of genesets; adjusted *p*-values were computed using Benjamini-Hochberg method [12], implemented in **p.adjust** of **R**.

### Neural stem cell differentiation

Lesch-Nyhan Disease hints interesting links between aberrant purine biosynthesis and neurophysiological and neurobehavioral disorder [19]. Although overall mechanisms are yet to be understood, previous studies discovered that mutations of HPRT (hypoxanthine guanine phosphoribosyltransferase) causes defects in neuronal growth and differentiation [20]. Here we focused on changes of network modules/pathways. We used time-series case versus control expression matrices *X* and *Y* constructed from the RNA-sequencing dataset (GSE42662) of 14 days of Dopaminergic (DA) neuronal differentiation of spherical neural masses (SNMs) [21]. The dataset includes 10 snapshots of days 0, 1, 2, 3, 4, 6, 8, 10, 12 and 14, generated by human DA differentiation protocol [22], where they characterize 14 days as three distinctive phases: (1) neuronal induction from days 0 to 4; (2) DA neuron induction from days 4 to 8; (3) DA neuron maturation from days 8 to 14.

#### Significant network modules

##### Network clustering

We ran our hierarchical network clustering algorithm, improved upon our previous work [23], on BioGRID physical and Reactome co-reaction networks separately, and resolved 173 of physical and 144 of co-reaction modules (see Methods for details). To ensure high quality of modules, we compared results with other network clustering methods. In extensive benchmark and cross-validation experiments, we found our algorithm substantially outperformed than others (see supplementary results; Fig.7 and 8).

##### Time-specific network modules changing the fate of stem cell differentiation

We focus on results of TED-dpm (TED and DPM-based geneset refinement) because of substantial improvement of statistical power and homogeneity of modules. As expected from the result of simulation study (Fig.2), TED-dpm was able to identify more significant modules than other methods. Geneset refinement steps effectively separated out noisy, or weak, gene expressions from genesets, highlighted true signals embedded in the sets. We confirmed that after the refinement other methods such as GSA significantly better than before (see supplementary Table.1, 2, 3 and 4). Fig.3 shows significantly changing modules found by TED-dpm method at familywise error rate 0.01. Due to space limit, we only show those non-trivially overlap with known canonical pathways [24], determined by hypergeometric test (FDR < 0.01 with overlap > 0.25).

**Figure 3.**
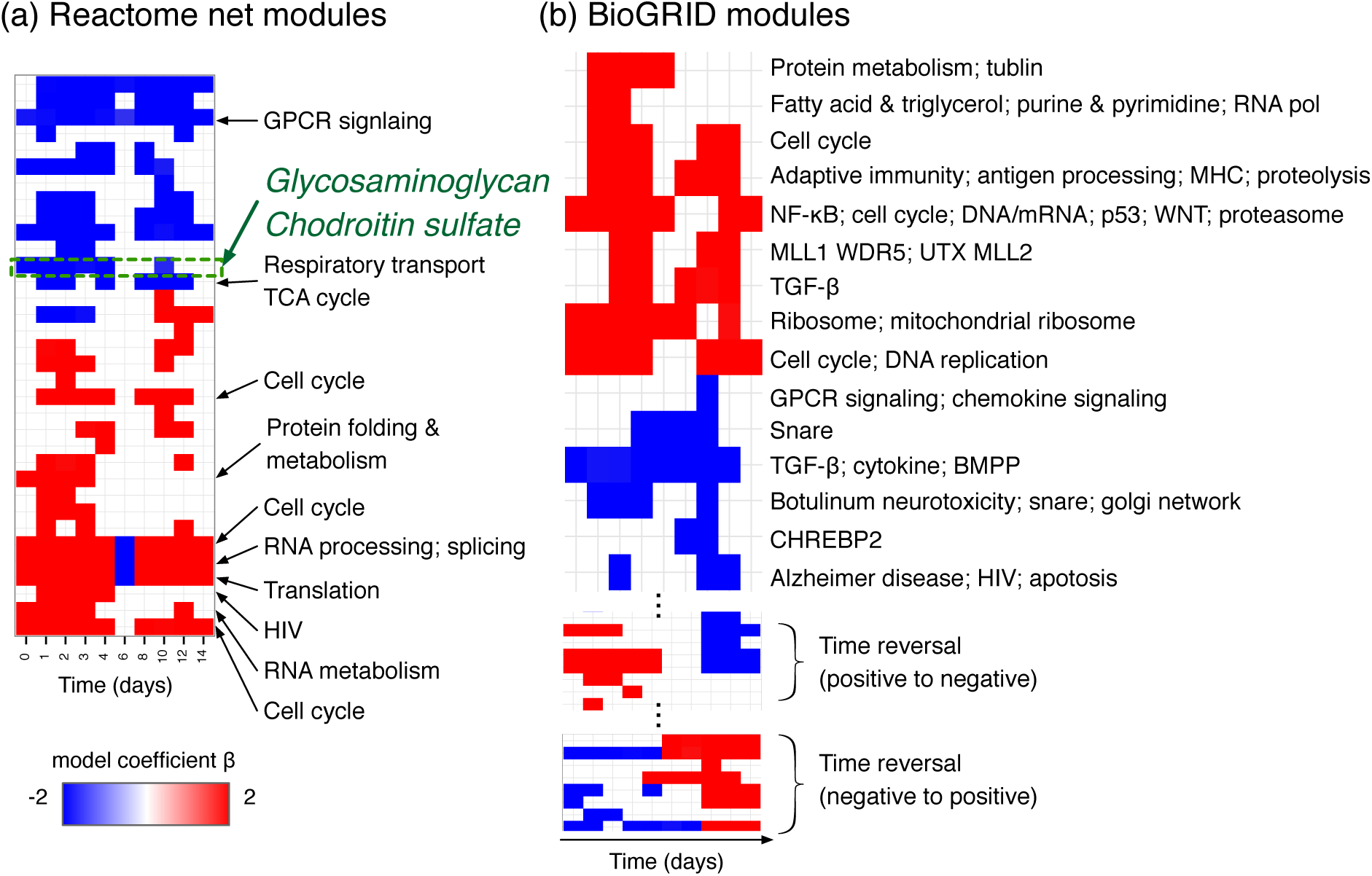
network moduels. We determined significance of these modules at familywise error rate (FWER) < 0.01; *p*-values corrected by Holm procedure (see e.g., [25]). Throughout all time course, modules related to cell cycle, neucleotide and downstream lipid metabolism were upregulated in control samples; in other words, activity of these modules was unusually dampened by HPRT gene KD. However, modules of GPCR signaling (Reactome), and a part of TGF- and pathways (BioGRID), were up-regulated in the KD cells. A large fraction of modules did not change direction of regulations either positive (red) or negative (blue). However, there are some modules undergo transient swapping (e.g., RNA processing in Reactome network, Fig.3a), or complete phase transition (e.g., anti-phasing modules in BioGRID network, Fig.3c). Unfortunately anti-phasing modules are poorly characterized in current knowledge; and they may have not been recognized as functional modules. On modules with almost equal amount of up- and down-regulated samples sample average of log2 ratio would only remain close to zero effect (row-wise summation).

##### Glycosaminoglycan complex

In Fig.4, we zoomed-in the glycosaminoglycan module of Reactome network. It is evident that tightly co-expressed core sub-complex (green dashed circle) plays a major role in the complex. The core includes glycan and chondroitin enzymes: B4GALT (*β*-1,4-galactosyltransferase), B3GAT (*β*-1,3-glucuronyltransferase), CHST (carbohydrate sulfotransferase), CHSY (chondroitin synthase), and CHPF (chondroitin polymerizing factor). In the knock-down HPRT cells, this core complex was initially up-regulated during neuronal induction phase until day 4, but down-regulated during DA neuron induction, then lost significant synchronous pattern during DA maturation (Fig.3a; Fig.4).

**Figure 4.**
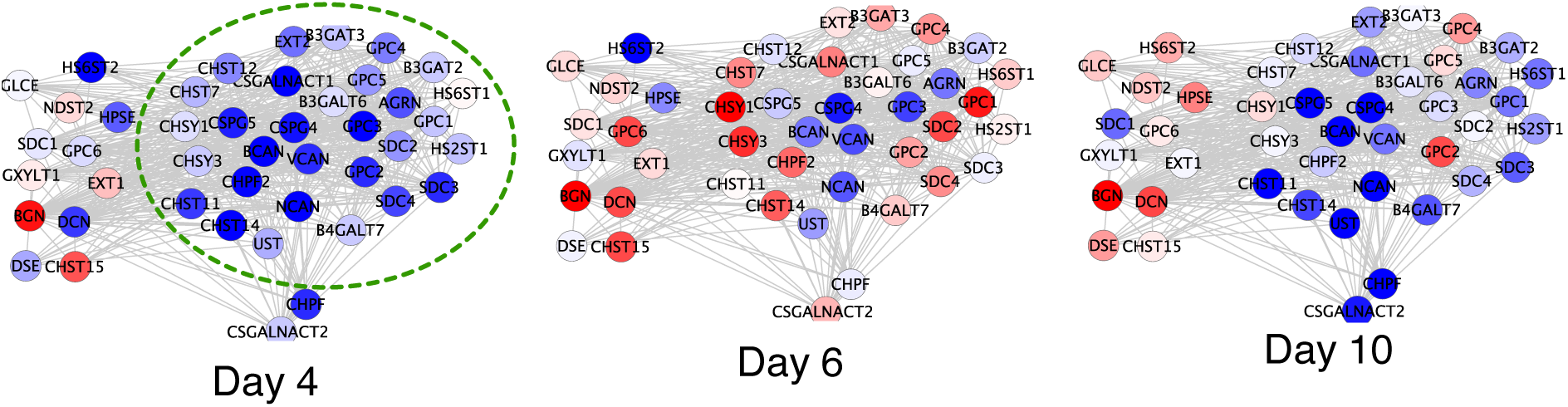
Glycosaminoglycan complex. Six sub-networks show transcriptomic dynamics of “Glycosaminoglycan” module in Reactome network. We show all parts of module inferred by network clustering; green dashed circle encloses parts included by TED-DPM. Nodes and edges represent genes and interactions of Reactome network (co-reaction). Genes were colored by expression level. Lowest log_2_(control/KD) ratio was colored blue (RGB 0,0,255); equal amount of control and KD colored white (RGB 255,255,255); highest colored red (RGB 255,0,0).

Along the same direction, recent studies show that neural stem cells can be identified specific glycan makrers, and many stem cell signaling pathways regulated and marked by post-transnational modifications of glycans attached to membrane [26]. Moreover, there was a systematic assay that mRNA level of many glycan enzymes strongly correlates with cell fate change of mouse embryonic stem cells [27]. In some cases, abnormal expression B4GALT family genes has been shown to promote multi-drug resistance in leukemia cells [28]. In sum, we may suggest an interesting hypothesis of LND: upon HPRT knockdown, cells experience dysfunctional purine metabolism, which directly leads to inappropriate exposition of glycans to cell surface; interplay of other signaling pathways (e.g., Wnt and GPCR) completely change cell fates departing from DA neurons. However, we may find better picture by follow-up experimental studies.

### Common signature of regulation

It was our surprise that a substantial faction of network modules were largely heterogeneous mixtures of TED models. In fact, we modified TED to address this nature. Although our network clustering algorithm recapitulated high-probability modules, we further tested to show heterogeneity of modules is indeed algorithm-independent.

#### Pathways are mixed

To answer that question, we incorporated TED models in Latent Dirichlet Allocation (LDA) framework [11]. One may think of LDA as probabilistic principal components. We looked for commonly observed expression patterns across genesets, and estimated mixing proportion of genesets. Common patterns are termd topics, which are TED topics, characterized by *β* parameters. We collected manually curated ≈ 1, 400 genesets (**c2.cp**) from MSigDB [24], which provides extensive coverage of known and manually curated pathways collected from various databases. On top of that, we also added Reactome and Biogrid network modules found in our network clustering.

We were able to cofirm that trained TED topics (Fig.5a,b) summarized what we repeatedly observed pattern of gene expressions in many genesets and network modules. Models either with 3 or 6 topics showed highest predictive performance in 10-fold cross validation experiments (see Supplementary result; Fig.6). But we mapped genesets and network modules to 3-TED topic space for simplified visualization (Fig.5c-e). Genesets are apparently mixed. Although many of genesets locate nearby the top corners of triangles (topic 2 in Fig.5a), a remaining portion of genesets are mixture of topics 0 and 1. Notably MIPS complexes are biased toward topic 2, which means that these complexes were overexpressed in control cells. It appears that MIPS complexes were established on normal cells. Therefore, future analysis based solely on MIPSs would have to be limited and may not discover new disease-specific pathways and complexes.

**Figure 5.**
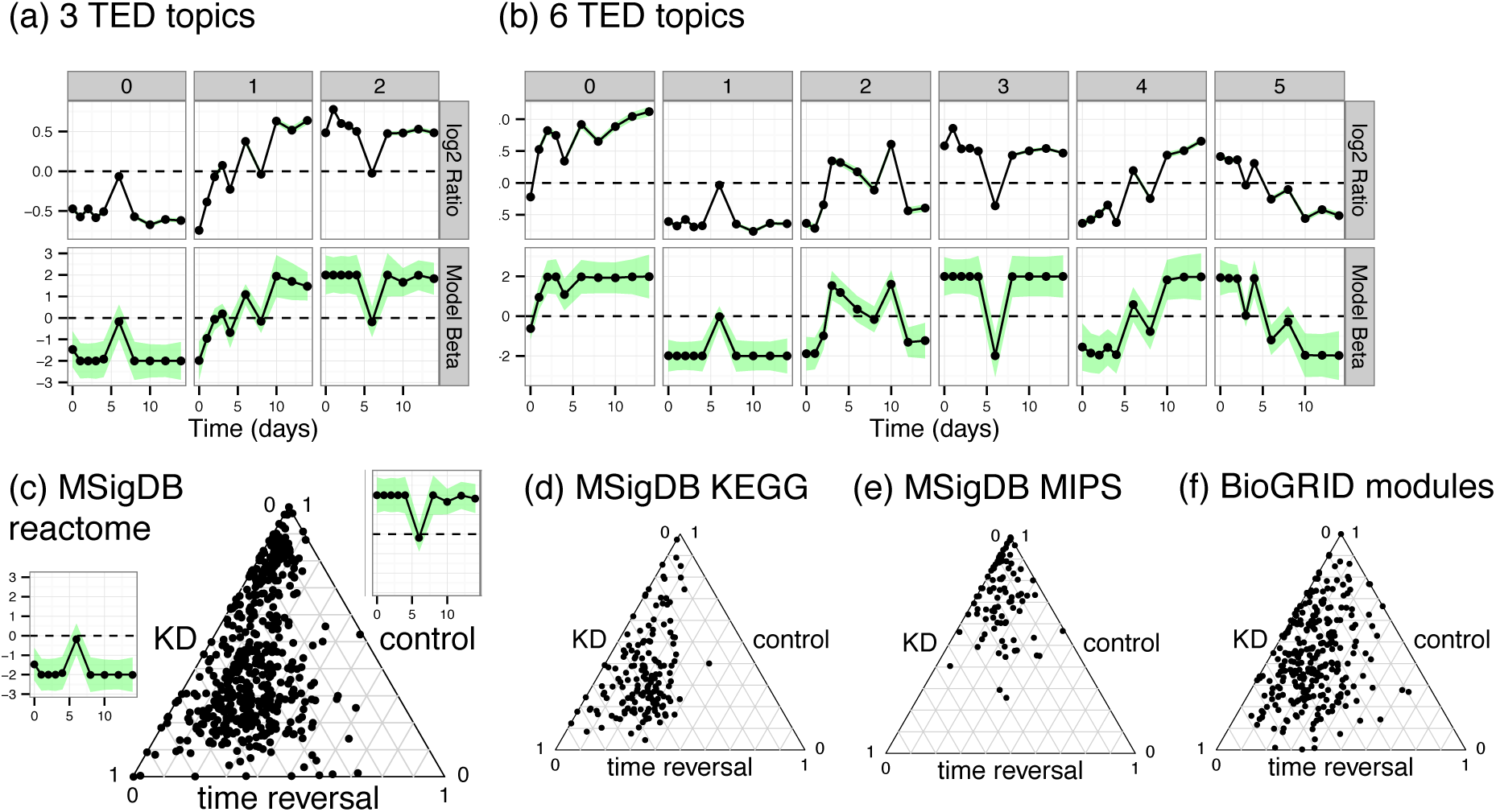
Genesets are mixed. (a) 3 distinctive TED topics. Each column shows each topic. Top row enumerates topic-specific average of *log*_2_(control/KD). Bottom row shows topic-specific model parameters *β* with 95% confidence bands (shaded area). (b) 6 TED topics. Top and bottom rows show 6 topic-specific data averages and model parameters. (c) Triplot of topic proportion. Each dot corresponds to MSigDB’s “reactome” genesets. Three vertices of simplex correspond to topics 0 to 2 of 3 TED topics (counterclockwise from the left corner of triangle). (d) Topic proportion of MIPS complexes (e) Topic proportion of BioGRID network modules.

**Figure 6.**
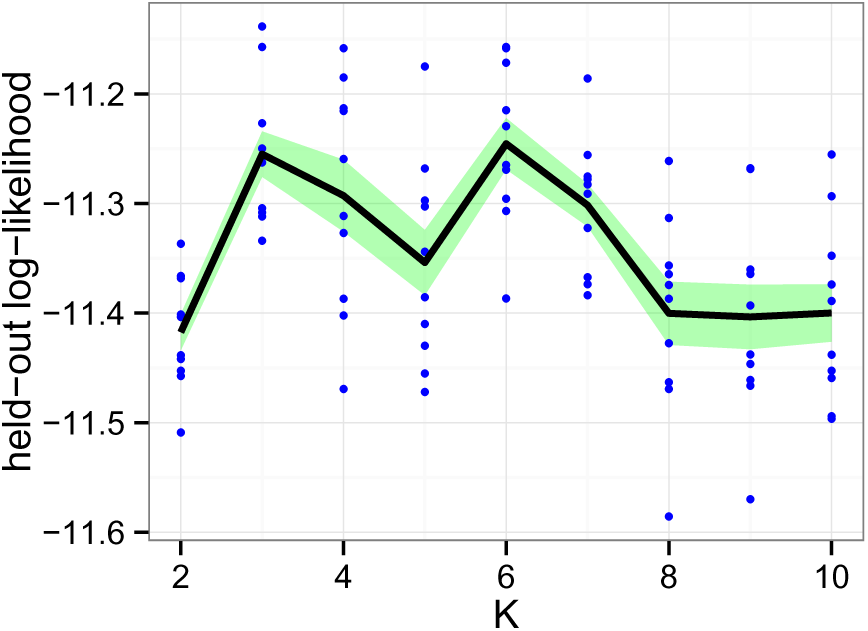
Bold the first sentence. Rest of figure 2 caption. Caption should be left justified, as specified by the options to the caption package.

**Figure 7.**
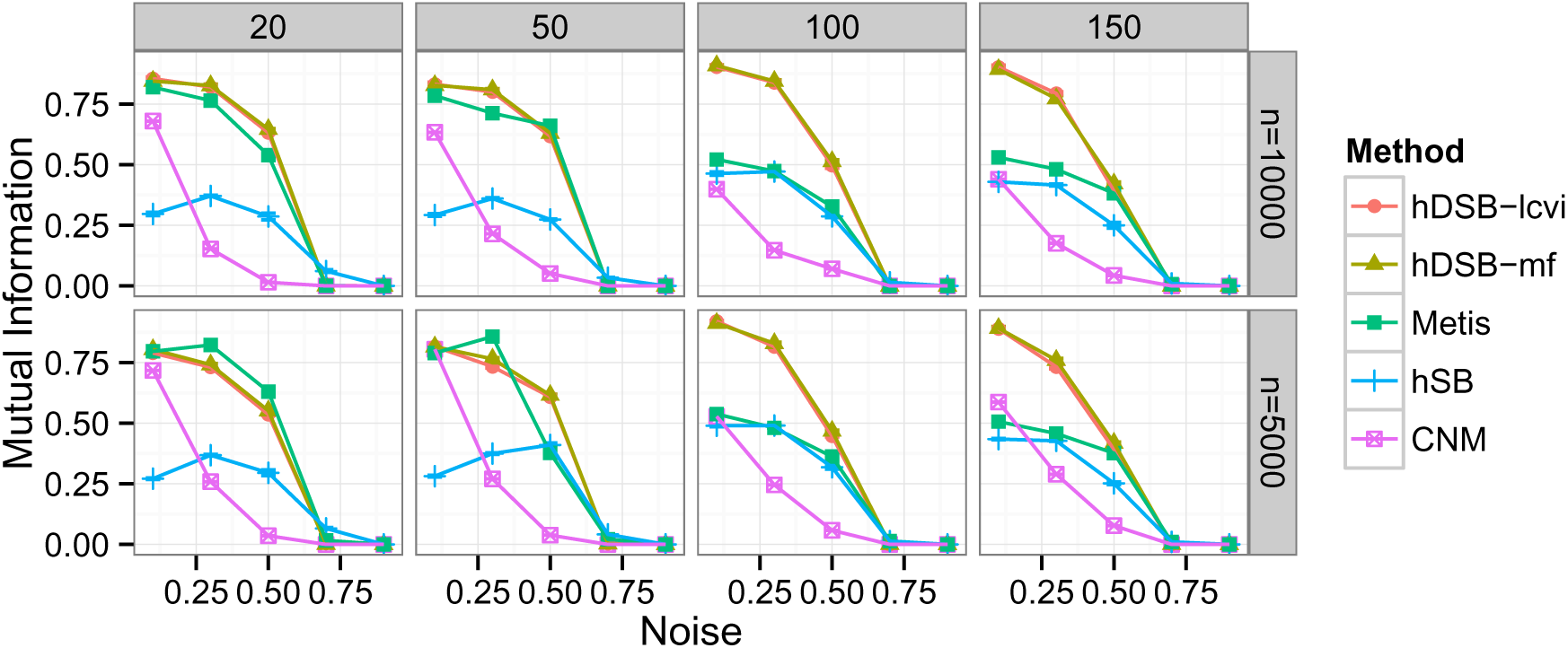
Bold the first sentence. Rest of figure 2 caption. Caption should be left justified, as specified by the options to the caption package.

**Figure 8.**
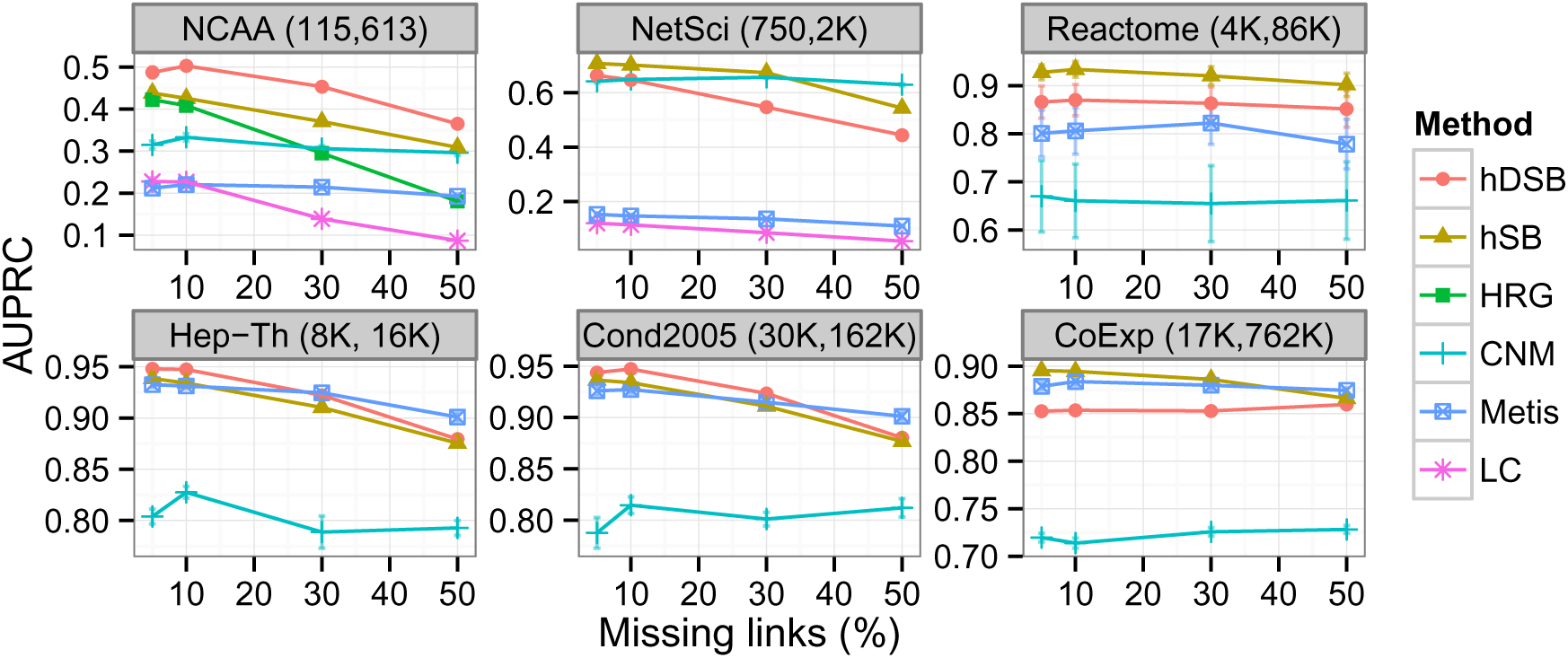
Bold the first sentence. Rest of figure 2 caption. Caption should be left justified, as specified by the options to the caption package.

## Conclusion

We developed a novel set-based method to annotate pre-defined genesets or modules inferred from networks. The methods were not only statistically more powerful, but also able to capture transient signals reliably. We designed TED and variants in two philosophy: classification is easier than density estimation (e.g., [29]). However, previous works approach the problem to sift out significant sets by unusual density of observations. We also wanted to do discriminative learning, which is usually easier than generative model fitting (e.g., [30]). TED and TED-variants outperformed in both synthetic and real-world than other geneset analysis methods. Moreover, we empirically demonstrated prevalent heterogeneity during the neural differentiation process. We also showed anti-phasing modules yet to be characterized by subsequent experiments and analysis.

## Methods

### Data preparation

#### Interactome

We constructed a physical interaction network dataset of 14, 995 proteins and 140, 006 interactions from BioGRID 3.1.94 [31], which are edges labeled physical. We also constructed a co-reaction network dataset of 4, 527 genes and 87, 947 interactions. We downloaded Reactome network database [32] from http://www.reactome.org/download/current/homo_sapiens.interactions.txt.gz. We used “current version” as of Aug 4 2013. We only included edges labeled reaction or neighbouring reaction, so as to identify modules, which could not be found in physical interaction networks.

#### Transcriptome

Over 98% of short reads were mapped to NCBI mm9 transcripts using tophat [33], counted by **htseq-count** (http://www-huber.embl.de/users/anders/HTSeq/doc/count.html) and normalized by **DESeq** taking into accounts of effective size factors [34]. The data was collected on mouse embryonic stem cells but mouse interactomes were rather poorly characterized compared to human. We mapped mouse genes to human genes using **biomaRt** [35]. Resulting *X* and *Y* matrices contained 10, 170 genes of 10 snapshots (days 0, 1, 2, 3, 4, 6, 8, 10, 12 and 14).

### Variational inference of TED model

#### Bayesian inference and hypothesis testing

We performed Bayesian inference on unknown parameters, especially the *β* parameters. We defined likelihood of observed *X, Y* under the parameter *β* of TED model (Eq.2), and the *β* parameters directly translate to significance of observed *X, Y*, quantifying steepness of decision boundary. Since we have 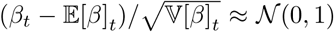, both approximately and asymptotically, we may construct level *α* Wald test of testing *H*_0_ : *β_t_* = 0 versus *H*_0_ : *β_t_* ≠ 0 as:

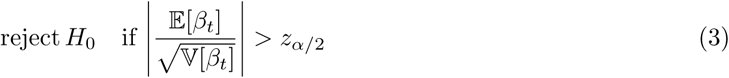

where *z*_*α*/2_ = Φ^−1^(1 − *α*/2). Then,

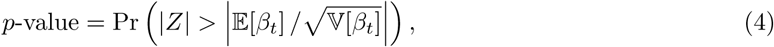

where *Z ~ N* (0, 1). In a strict sense, our hypothesis testing is more akin to controlling false discovery rate rather than Type I error. We may consider TED as a “discriminative” version of Sun and Cai [36,37].

#### Bayesian sparse prior

Now, let us discuss how to estimate 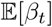 and 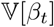 taking into account of temporal smoothness. We introduced the Fused Lasso prior [38] on to *β* parameters.

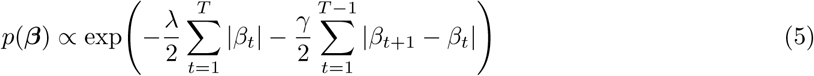

where *λ* and *γ* control static and kinetic sparsity of parameters. Under appropriate *λ* and *γ* we capture sparse and temporally smooth *β*. However, estimation of *β* under the Fused Lasso prior may not be technically difficult unlike regular Lasso prior [39]. Instead of direct usage, we used equivalent Bayesian formulation to handle sparsity more gently (see [40] and [41] for details and proofs).

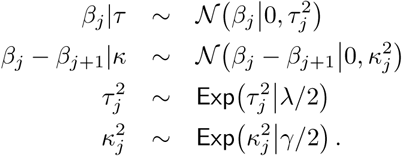

#### Non-conjugate variational inference

To work around non-conjugate relationship between the likelihood function (Eq.2) and prior (Eq.5), we estimated posterior probability by the non-conjugate variational inference [42]. First we find optimal *β* by coordinate descent algorithm [43], then construct approximate distribution *q*(*β*) by the Laplace method [42].

##### Distribution of β

First of all, it is not hard to verify convexity of negative log-likelihood function:

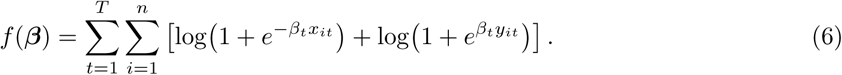

Like other GLMs, we can construct local quadratic approximation [43, 44]. Using these results, we can construct the following local quadratic form *g*(*β*) ≈ *f* (*β*) at current estimate 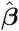 defined by

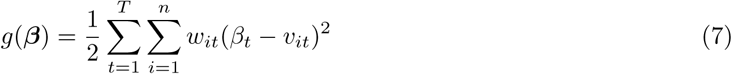

where

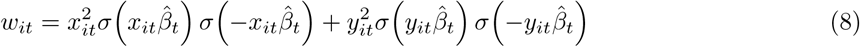

and

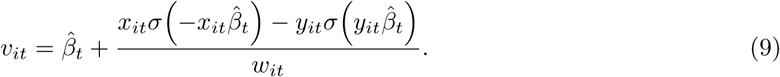

With the prior on *β*, we find the solution by solving

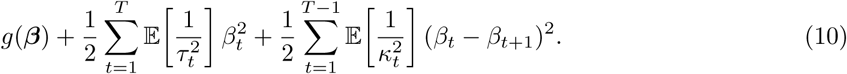

For each *t* ∈ {2, *T* − 1}, we can iteratively update *β_t_* until convergence as follows.

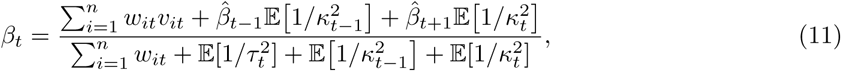

we drop the *t* − 1 term for *t* = 1 and *t* + 1 for *t* = *T*.

Following Wang and Blei [42], we have

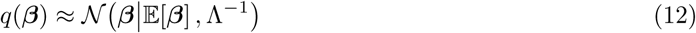

where 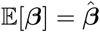 and tridiagonal precision matrix

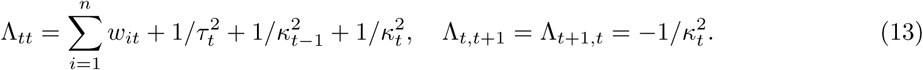

For subsequent updates, we need covariance matrix Σ ≡ Λ^*−*1^. Exploiting special structure of Λ, we can easily resolve tri-diagonal elements of Σ, i.e., Σ_*tt*_ and Σ_*t,t*+1_ by linear time algorithms (e.g., [45] or Appendix B of [46]).

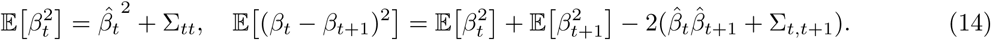

##### Distribution of τ, κ

From the general result of Kyung *et al.* [41], although tedious derivations yield the same results, we have 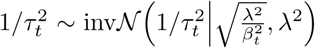 and 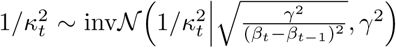. Therefore, we can find mean-field solution required for the update of *q*(*β*) and empirical Bayes estimation penalty terms as:

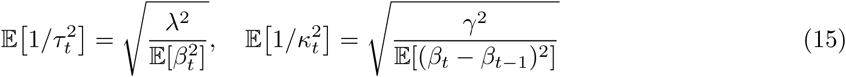

and

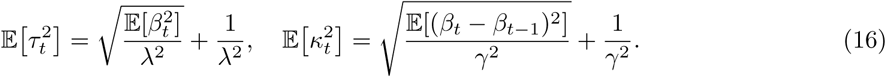

#### Empirical Bayes

We adjust a proper degree of penalties, *λ* and *γ*, by optimizing the marginal likelihood weighted by the variational distributions over *β* and *τ*, *κ* (see [40, 41] for details). We update *λ* and *γ* as:

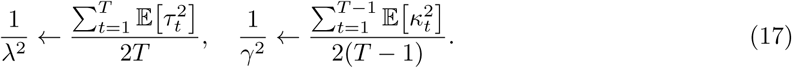

until convergence.

#### Overall inference algorithm

Initially we set 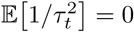 for *t* ∈ [*T*] and 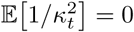 for *t* ∈ [*T* − 1]. Then, we repeat the following steps until convergence:

1. Optimize 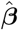 (Eq.11) and estimate 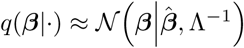 (Eq. 12).
2. Calculate covariance matrix Σ ≡ Λ^*−*1^ and 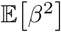 and 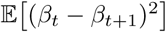 (Eq. 14).
3. Update 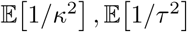 (Eq. 15).
4. Alternate update of 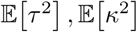 (Eq.16) and empirical Bayes estimation of *λ*, *γ* until convergence (Eq.17).

C++ implementation of TED and related methods are available in public repository (https://code.google.com/p/glmblock).

### Mixture and admixture of TED models

#### Locally collapsed latent update

So far variational inference algorithm assumed a single TED model. For both mixture or admixture models, we need a scoring function to evaluate probabilistic assignment of each object, i.e., a pair of expression vectors x_*i*_ and y_*i*_, to TED models.

We found “locally collapsed” variational inference (LCVI) [47] works better than regular mean-field approximation methods. Suppose for each TED model *k* we have estimated variational *q_k_*(*β*). Essentially LCVI helps avoid bad local optima infusing adequate stochasticity. Given them, latent assignment of x and y is conditionally independent. Let *c_i_* ∈ [*K*] be a random variable of *i*’s model assignment, although sometimes *K* → ∞. We integrate out uncertainty of with respect *q_k_* 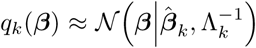 previously determined by each *k*-th model

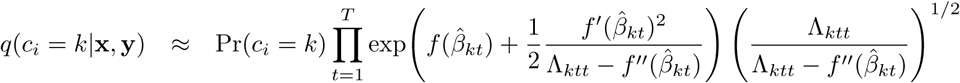

where *β_kt_* denotes *t*-th element of *β_k_* vector; Λ_*ktt*_ denotes (*t, t*) element of Λ_*k*_ matrix. We calculate appropriate prior factor Pr(*c_i_* = *k*) depending on DPM or LDA.

But we may not have previous observation at all. For instance, in DPM inference, we can discover a new model which was never explored. We simply treat *β_t_* ~ *N* (*β_t_*|0, *τ*^2^) with 1/*τ*^2^ = ∞; we use the fact that 1/*τ*^2^ ≈ *λ*/|*β_t_*| → ∞ as *β* → 0. We elaborated detailed derivations and computationally e cient update strategies in the appendix.

#### Dirichlet Process Mxiture

Here we briefly mention sketch of overall algorithm. First we randomly assign expression pairs (x_*i*_, y_*i*_) to models and update models given data. Iteratively we sample assignments of (x_*i*_, y_*i*_), which works like E-step of Expectation Maximization [48]; then, we perform variational update of each TED model including empirical Bayes routines, and this works like M-step. Interested readers may refer to variational inference methods of general DPMs (e.g., [47, 49]). Our algorithm was essentially equivalent to Wang and Blei [47].

#### Latent Dirichlet Allocation (admixture model)

LDA [11] models corpus of documents, while assigning topics to words and topic proportions to documents. Each topic is defined by its own topic-specific word frequency vector. Here, we treat genesets as documents and genes as words. Topics are defined by TED models. We sampled each gene’s topic assignment according to LCVI score (Eq.18). Each TED model was trained by genes assigned in online stochastic learning [50]. While gradually observing genesets, we udpated TED topics. We tested both batch and online learning, but online learning worked in a more scalable way and resolved equally likely models. Again, details can be found in the original paper [11] and algorithm paper [50].

### Network clustering

We extended our previous method, called dynamic hierarchical model (DyHM) [23] in two aspects: (1) we updated base-model to take into accounts of empirical degree distribution [51]; (2) we sped up the overall algorithm using dynamic programming technique. Source codes are available in public repository (https://code.google.com/p/hsblock).

**Table 1.**
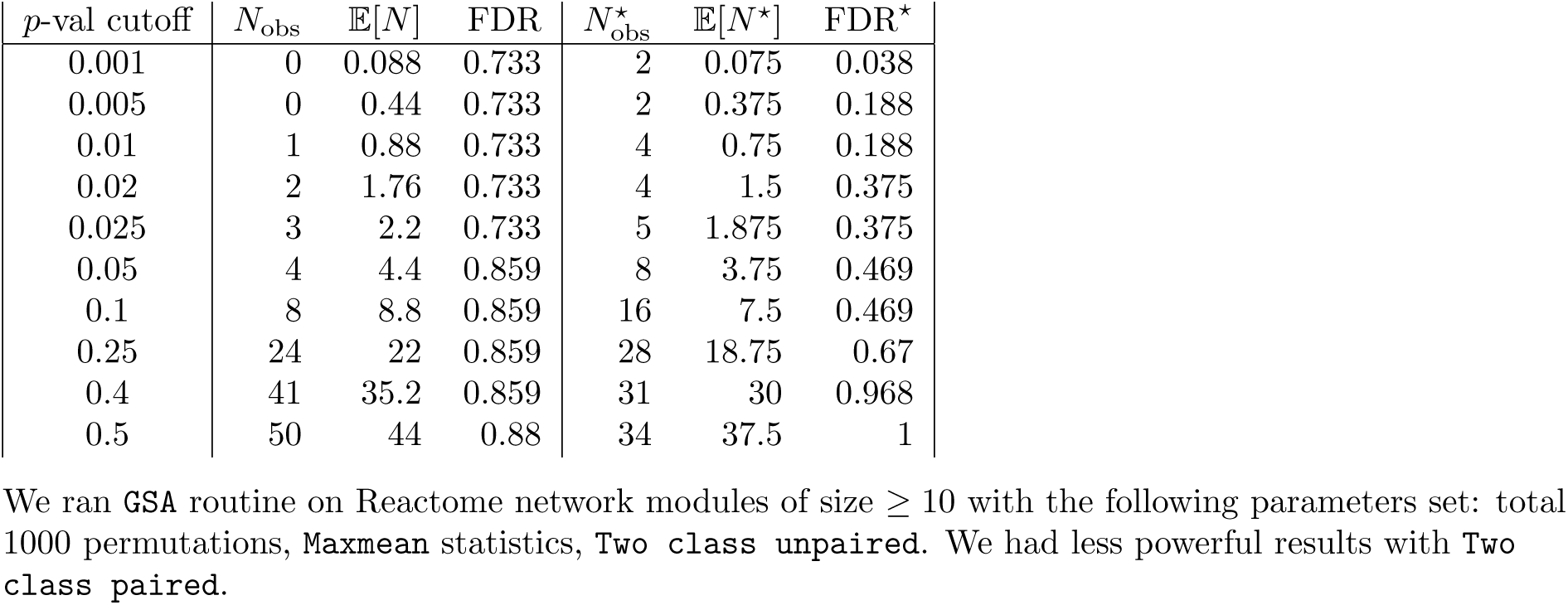
GSA results testing up-regulation (hi) of Reactome network modules

| $p$-val cutoff | $N_{\text{obs}}$ | $\mathbb{E}[N]$ | FDR | $N_{\text{obs}}^*$ | $\mathbb{E}[N^*]$ | FDR* |
| --- | --- | --- | --- | --- | --- | --- |
| 0.001 | 0 | 0.088 | 0.733 | 2 | 0.075 | 0.038 |
| 0.005 | 0 | 0.44 | 0.733 | 2 | 0.375 | 0.188 |
| 0.01 | 1 | 0.88 | 0.733 | 4 | 0.75 | 0.188 |
| 0.02 | 2 | 1.76 | 0.733 | 4 | 1.5 | 0.375 |
| 0.025 | 3 | 2.2 | 0.733 | 5 | 1.875 | 0.375 |
| 0.05 | 4 | 4.4 | 0.859 | 8 | 3.75 | 0.469 |
| 0.1 | 8 | 8.8 | 0.859 | 16 | 7.5 | 0.469 |
| 0.25 | 24 | 22 | 0.859 | 28 | 18.75 | 0.67 |
| 0.4 | 41 | 35.2 | 0.859 | 31 | 30 | 0.968 |
| 0.5 | 50 | 44 | 0.88 | 34 | 37.5 | 1 |

We ran **GSA** routine on Reactome network modules of size $\geq 10$ with the following parameters set: total 1000 permutations, **Maxmean** statistics, **Two class unpaired**. We had less powerful results with **Two class paired**.

**Table 2.** GSA results testing down-regulation (lo) of Reactome network modules

| $p$-val cutoff | $N_{\text{obs}}$ | $\mathbb{E}[N]$ | FDR | $N_{\text{obs}}^*$ | $\mathbb{E}[N^*]$ | FDR* |
| --- | --- | --- | --- | --- | --- | --- |
| 0.001 | 0 | 0.088 | 0.819 | 1 | 0.075 | 0.075 |
| 0.005 | 0 | 0.44 | 0.819 | 2 | 0.375 | 0.188 |
| 0.01 | 0 | 0.88 | 0.819 | 4 | 0.75 | 0.188 |
| 0.02 | 2 | 1.76 | 0.819 | 4 | 1.5 | 0.375 |
| 0.025 | 2 | 2.2 | 0.819 | 5 | 1.875 | 0.375 |
| 0.05 | 4 | 4.4 | 0.819 | 9 | 3.75 | 0.417 |
| 0.1 | 9 | 8.8 | 0.819 | 17 | 7.5 | 0.441 |
| 0.25 | 26 | 22 | 0.819 | 28 | 18.75 | 0.67 |
| 0.4 | 43 | 35.2 | 0.819 | 32 | 30 | 0.938 |
| 0.5 | 49 | 44 | 0.898 | 34 | 37.5 | 1 |

We ran **GSA** routine on Reactome network modules of size $\geq 10$ with the following parameters set: total 1000 permutations, **Maxmean** statistics, **Two class paired**. We had less powerful results with **Two class unpaired**.

**Table 3.** GSA results testing up-regulation (hi) of BioGRID network modules

| $p$-val cutoff | $N_{\text{obs}}$ | $\mathbb{E}[N]$ | FDR | $N_{\text{obs}}^*$ | $\mathbb{E}[N^*]$ | FDR* |
| --- | --- | --- | --- | --- | --- | --- |
| 0.001 | 0 | 0.085 | 0.85 | 5 | 0.204 | 0.041 |
| 0.005 | 0 | 0.425 | 0.85 | 5 | 1.02 | 0.204 |
| 0.01 | 0 | 0.85 | 0.85 | 5 | 2.04 | 0.34 |
| 0.02 | 2 | 1.7 | 0.85 | 12 | 4.08 | 0.34 |
| 0.025 | 2 | 2.125 | 0.944 | 13 | 5.1 | 0.392 |
| 0.05 | 4 | 4.25 | 0.944 | 22 | 10.2 | 0.434 |
| 0.1 | 9 | 8.5 | 0.944 | 47 | 20.4 | 0.434 |
| 0.25 | 19 | 21.25 | 1 | 73 | 51 | 0.699 |
| 0.4 | 29 | 34 | 1 | 84 | 81.6 | 0.971 |
| 0.5 | 35 | 42.5 | 1 | 92 | 102 | 1 |

We ran **GSA** routine on BioGRID network modules of size $\geq 10$ with the following parameters set: total 1000 permutations, **Maxmean** statistics, **Two class unpaired**. We had less powerful results with **Two class paired**.

**Table 4.**
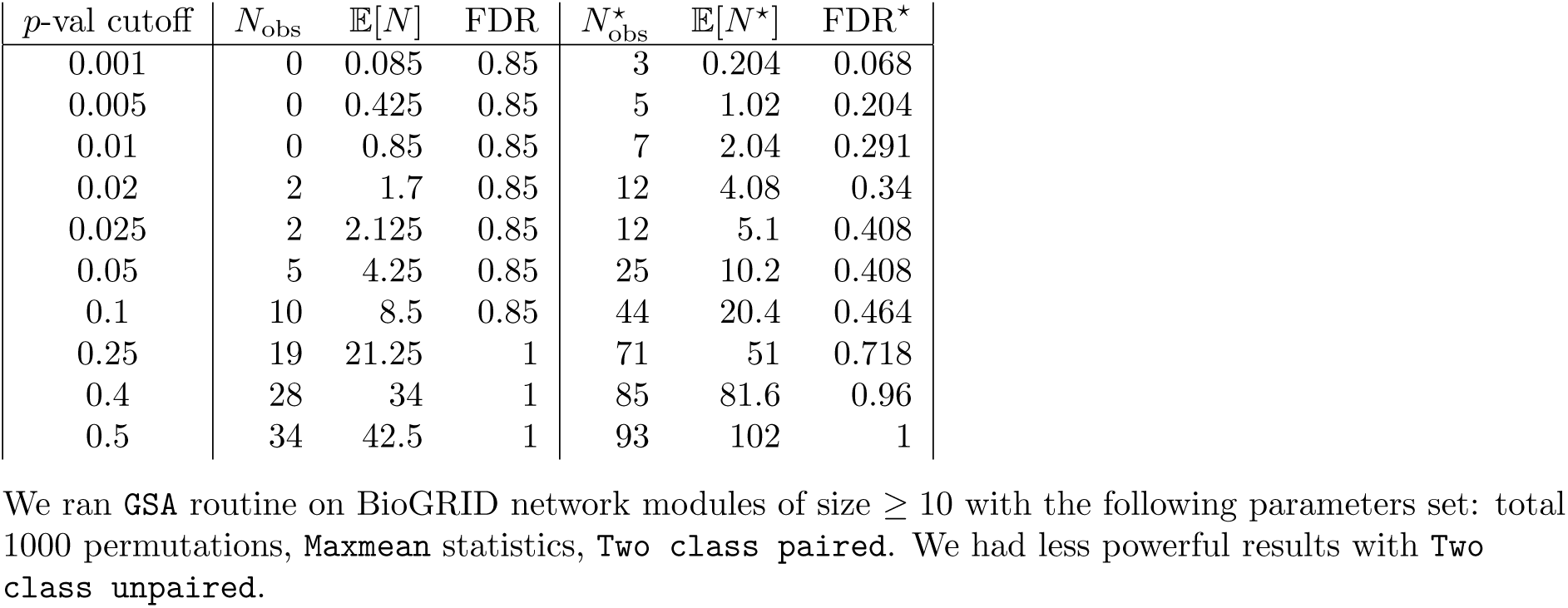
GSA results testing down-regulation (lo) of BioGRID network modules

## Appendix: Derivation of update equations

### TED

#### Locally Collapsed Variational Inference

Again, to circumvent non-conjugate relations, we approximate the likelihood by second order Taylor expansion,

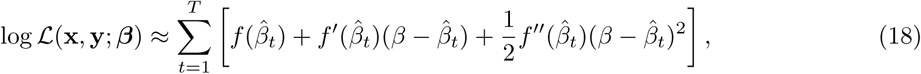

where 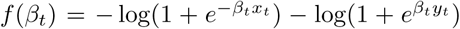. Then, posterior probability of (x_*i*_; y_*i*_) assignment to model *k* is straightforward.

For simplicity, let 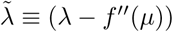 and 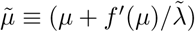

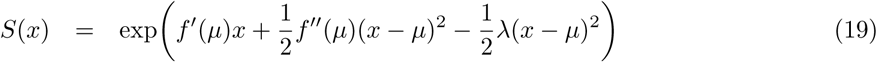

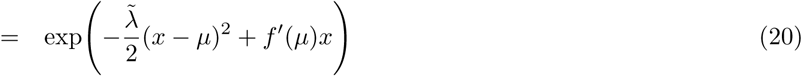

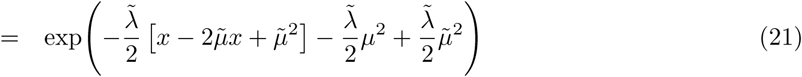

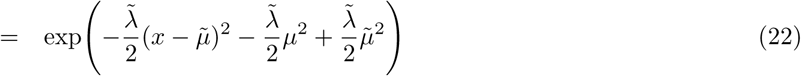

Then,

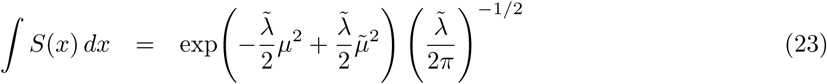

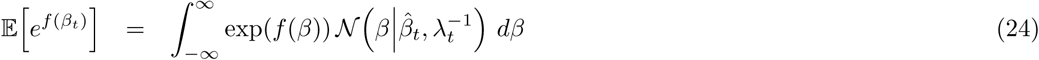

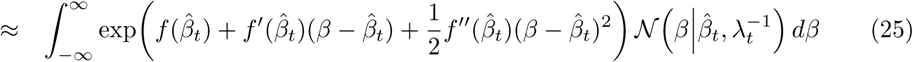

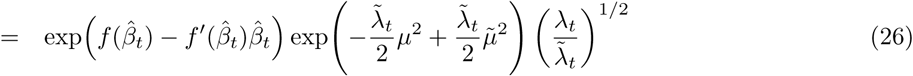

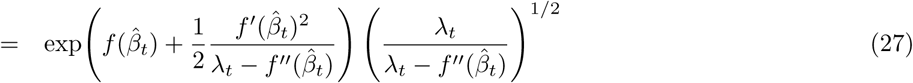

## Appendix: supplementary results

### Latent Dirichlet Allocation

10-fold cross validation

### Geneset analysis (GSA)

We ran GSA [14]

### Network clustering methods

